# Altered fibrin clot structure contributes to thrombosis risk in severe COVID-19

**DOI:** 10.1101/2021.09.17.460777

**Authors:** Malgorzata Wygrecka, Anna Birnhuber, Benjamin Seeliger, Laura Michalick, Oleg Pak, Astrid-Solveig Schultz, Fabian Schramm, Martin Zacharias, Gregor Gorkiewicz, Sascha David, Tobias Welte, Julius J. Schmidt, Norbert Weissmann, Ralph T. Schermuly, Guillermo Barreto, Liliana Schaefer, Philipp Markart, Markus C. Brack, Stefan Hippenstiel, Florian Kurth, Leif E. Sander, Martin Witzenrath, Wolfgang M. Kuebler, Grazyna Kwapiszewska, Klaus T. Preissner

## Abstract

The high incidence of thrombotic events suggests a possible role of the contact system pathway in COVID-19 pathology. Here, we demonstrate altered levels of factor XII (FXII) and its activation products in two independent cohorts of critically ill COVID-19 patients in comparison to patients suffering from severe acute respiratory distress syndrome due to influenza virus (ARDS-influenza). Compatible with this data, we report rapid consumption of FXII in COVID-19, but not in ARDS-influenza, plasma. Interestingly, the kaolin clotting time was not prolonged in COVID-19 as compared to ARDS-influenza. Using confocal and electron microscopy, we show that increased FXII activation rate, in conjunction with elevated fibrinogen levels, triggers formation of fibrinolysis-resistant, compact clots with thin fibers and small pores in COVID-19. Accordingly, we observed clot lysis in 30% of COVID-19 patients and 84% of ARDS-influenza subjects. Analysis of lung tissue sections revealed wide-spread extra- and intra-vascular compact fibrin deposits in COVID-19. Together, our results indicate that elevated fibrinogen levels and increased FXII activation rate promote thrombosis and thrombolysis resistance *via* enhanced thrombus formation and stability in COVID-19.

## Introduction

Severe acute respiratory syndrome coronavirus 2 (SARS-CoV2) is a corona virus that causes a multisystem disease emanating from the respiratory tract designated as a coronavirus disease (COVID)-19^1–3^. Rapidly accumulating data suggests that a major underlying molecular mechanism in COVID-19-related morbidity and mortality is widespread endothelial injury associated with hyperactivation of the immune system, consequently leading to numerous haemostasis abnormalities^4–6^. Accordingly, next to markedly elevated levels of pro- and anti-inflammatory mediators such as interleukin (IL)-6, IL-2R, IL-10, and tumor necrosis factor-α (TNF-α), elevated levels of D-dimer, fibrinogen, and prolonged prothrombin time (PT) have been reported in severely ill COVID-19 patients^7–10^. The clinical relevance of these processes is highlighted by the association between abnormal levels of D-dimer and the 28-day mortality in patients with COVID-19^11–15^, and post-mortem studies stressing the presence of micro-thrombi and capillarostasis in the lungs of affected subjects^16,17^.

The high incidence of thrombotic events, in particular deep vein thrombosis and pulmonary embolism, in conjunction with mildly prolonged activated partial thromboplastin time (APTT)^18,19^, suggests a possible role of coagulation factor XII (FXII) in COVID-19 coagulopathy. FXII is a serine protease of the contact-phase system of blood coagulation and circulates in plasma as a single-chain zymogen^20^. Following contact with anionic surfaces such as kaolin, but also extracellular RNA (eRNA) released from damaged cells^21^, neutrophil extracellular traps (NETs)^22^, or polyphosphates secreted from activated platelets^23^, FXII undergoes autoactivation to αFXIIa (herein referred to as FXIIa)^24^. FXIIa cleaves plasma prekallikrein (PK) to kallikrein (PKa), which in turn reciprocally activates FXII and amplifies FXIIa generation^25^. As a consequence, the plasma kallikrein-kinin system is activated, leading to the release of the vasodilatory and vascular barrier disrupting peptide bradykinin (BK) from high molecular weight kininogen (HK)^26,27^. Overall, activation of the contact-phase system contributes to an increased production of thrombin and fibrin, although FXIIa/PKa-mediated conversion of plasminogen to plasmin may have a minor effect on fibrinolysis.

A congenital deficiency of FXII in humans does not cause any bleeding complications, suggesting that FXII is dispensable for physiological haemostasis and fibrin formation^28^. However, the contact phase pathway may play an important role in thrombosis development when contact surfaces are exposed in scenarios such as trauma injury or bacterial and viral infections^29,30^. Indeed, numerous *in vivo* studies have confirmed a critical function of FXII in thrombus growth and stabilization under the mentioned conditions and provided the rationale for the development of new FXIIa inhibitors, which ensure thrombo-protection in patients without causing a bleeding complications^29,31,32^.

Given the high incidence of thromboembolic complications in severely ill COVID-19 patients^18,19^, we investigated the contribution of FXII to fibrin formation and fibrinolysis in this patient cohort in comparison to patients infected with the influenza virus.

## Results

### FXII is activated in severely ill COVID-19 patients

In the discovery cohort, the plasma levels of FXII were decreased in severe COVID-19 patients as compared to controls (Figure 1A, B; moderate is defined by WHO severity score: 3–4, hospitalized, no invasive ventilation; severe is defined by WHO severity score: 5–7, high flow O_2_ or intubated and mechanically ventilated). Disappearance of FXII in plasma typically corresponds to its activation and conversion into two chain FXIIa protein composed of the 50 kDa-heavy chain and 30-kDa light chain. Detection of FXIIa plasma is, however, hindered by its raid inactivation and complex formation with C1 esterase inhibitor (C1INH). Thus to better monitor the presence of FXIIa in COVID-19 plasma, we measured products of its activation, such as HK and PKa. As expected, disappearance of FXII in plasma was accompanied by HK cleavage, seen as diminished signal intensity of intact HK band at 130 kDa (Figure 1C, D). A decrease in intact HK levels was associated with the appearance of cleaved HK fragments: the cleaved HK light chain band migrating at 55 kDa and an additional 45-kDa band representing a degradation product of 55-kDa cleaved HK light. To further examine whether the reduction in intact levels of FXII and HK is a result of contact system activation, we measured the activity of plasma PKa. PKa-like activity was markedly elevated in severe COVID-19 patients in comparison to donors and patients suffering from moderate SARS-CoV2 infection (Figure 1E). Furthermore, a strong negative correlation between the levels of intact HK and PKa-like activity in plasma of severe COVID-19 patients was observed (Figure 1F). Purified plasma proteins and deficient plasma samples were used to prove the specificity of the bands shown in western blots (Figure 1A, C; right panels).

**Figure 1.**
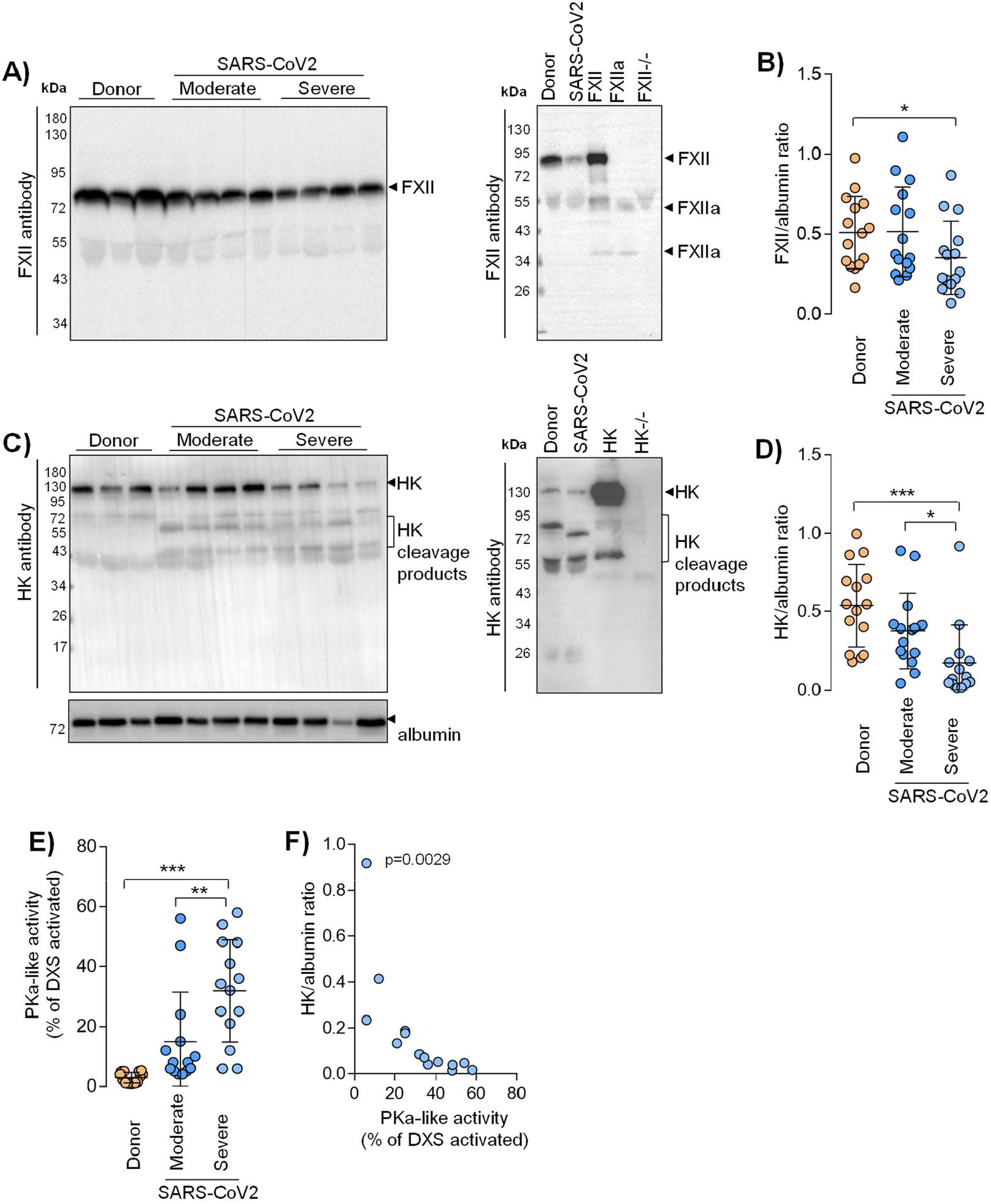
Activation of the contact phase system in plasma of critically ill COVID-19 patients. A,C) Western blot analysis (left panels) of factor XII (FXII) A) and high molecular weight kininogen (HK) C) in plasma from moderate and severe COVID-19 patients (infected with SARS-CoV2) and donors. Four out of 15 moderate and severe COVID-19 patients and 3 out of 15 donors are demonstrated. Rights panels show the specificity of the antibodies used. B, D) Densitometric analysis of A) and C), respectively. COVID-19 moderate/severe n=15, donor n=15. E) PKa-like activity in plasma from moderate (n=14) and severe (n=14) COVID-19 patients and donors (n=15). F) Correlation between the levels of intact HK and PKa-like activity in plasma of severe Covid-19 patients. n=14. Correlation is performed using Spearman’s rank correlation coefficient. *p<0.05, **p<0.01, ***p<0.001.Data in B), D), and E) are shown as mean+/-SD.

### Fibrinogen and FXIIa regulate fibrin network density in COVID-19

To assess whether enhanced activation of FXII in critically ill COVID-19 patients represents a characteristic feature of SARS-CoV-2 infection, we analyzed plasma samples of patients suffering from acute respiratory distress syndrome (ARDS) due to influenza virus infection. The decrease in FXII plasma levels in severe COVID-19 was confirmed in the validation cohort of the patients. Furthermore, the levels of FXII in COVID-19 were significantly lower than those in ARDS-influenza (Figure 2A). Surprisingly yet, the lag phase in fibrin formation, triggered by the FXII activator kaolin, was shorter in plasma of critically ill COVID-19 patients as compared to patients suffering from ARDS-influenza (Figure 2B). The phenomenon, which might be explained by markedly higher plasma levels of FVIII:C in COVID-19 than ARDS-influenza (Figure 2C). Notably, both COVID-19 and ARDS-influenza patients received the same daily dose of unfractionated heparin, excluding iatrogenic anticoagulation as a cause of prolonged kaolin-triggered clotting time in ARDS-influenza. In addition, we excluded lupus anticoagulant and the presence of anti-FXII antibodies as a cause of FXII deficiency in critically ill COVID-19 patients in our cohort (data not shown).

**Figure 2.**
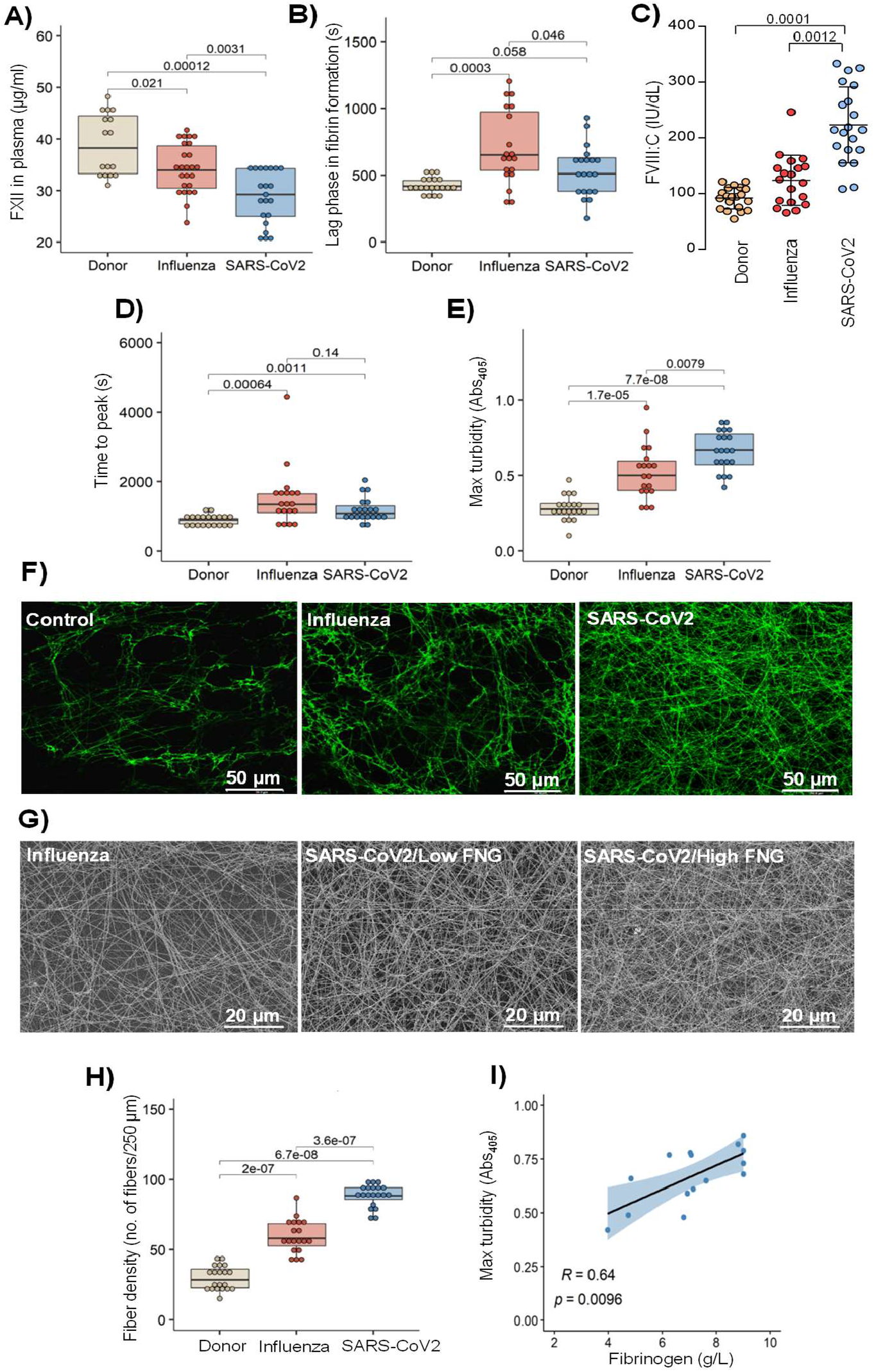
Dense fibrin clots are formed in severe COVID-19 plasma. A) Factor XII (FXII) levels in plasma of ARDS-influenza (Influenza; n=25) and severe COVID-19 (SARS-CoV2; n=21) patients as well as donors (n=16). B) Lag phase in fibrin formation-triggered by kaolin. Influenza, n=19; SARS-CoV2, n=20; donor, n=20. C) FVIII activity (FVIII:C) in patient and donor plasma. Influenza, n=19; SARS-CoV2, n=20; donor, n=20. Mean+/-SD is shown. D, E) Time to reach the turbidity peak D) and maximum (Max) turbidity E) values for Influenza (n=19), SARS-CoV2 (n=20) and donor (n=20) plasma. Clot formation was induced by the addition of kaolin to plasma. F) Laser scanning confocal microscopy images of fibrin fibers in clots formed from Influenza (n=19), SARS-CoV2 (n=20), and donor (n=20) plasma. Representative pictures are demonstrated. G) Scanning electron microscopy images of fibrin network in clots generated from Influenza as well as low- and high-fibrinogen (FNG) SARS-CoV2 plasma. Representative pictures are demonstrated. H) Fibrin fiber density in donor (n=20), ARDS-Influenza (n=19) and COVID-19 (n=20) clots. Per patient 3 separate clots were prepared, 5 pictures were taken in different areas of the clots and fibril density was determined in all pictures. I) Correlation between Max turbidity values and FNG levels in plasma of COVID-19 patients. SARS-CoV2-infected patients with available FNG levels are included into the analysis (n=15). Correlation is performed using Spearman’s rank correlation coefficient. Data in A), B), D), E), and H) are shown as single data points with boxplot overlay indicating median and interquartile range.

Further analysis of kaolin-triggered plasma clotting time revealed an increase in the time to reach the turbidity peak in both patient groups as compared to control, but no difference between ARDS-influenza and severe COVID-19 (Figure 2D). The density of the clot (indicated by the maximum turbidity measurement) was higher in both patient groups as opposed to control. A direct comparison between clots of ARDS-influenza and severe COVID-19 showed significantly higher maximal turbidity values in the latter group (Figure 2E). Visualization of fibrin clots by laser scanning confocal microscopy and scanning electron microscopy revealed an increase in fibrin structure compactness with thinner fibers and smaller pores in clots from COVID-19 plasma, as compared to clots generated in plasma obtained from ARDS-influenza patients (Figure 2F-H). A detailed analysis of the clots generated from plasma of severe COVID-19 patients demonstrated association between packing density of fibrin fibers and plasma fibrinogen concentration, with dense fibrin network in clots formed in plasma of patients exhibiting high fibrinogen levels (Figure 2G). Accordingly, a strong positive correlation between maximum turbidity values and fibrinogen concentration in plasma of critically ill COVID-19 patients was noted (Figure 2I).

As the architecture of fibrin clots may be influenced not only by fibrinogen (FNG) but also FXIIa^33,34^, we next analyzed the impact of these two proteins on the clot structure in a purified system. As depicted in figure 3A high concentrations of fibrinogen increased peak turbidity values and this effect was potentiated by the addition of FXIIa. Accordingly, corn trypsin inhibitor (CTI), the inhibitor of FXIIa, reduced maximum turbidity of the clot generated by mixing fibrinogen and FXIIa (Figure 3B).

**Figure 3.**
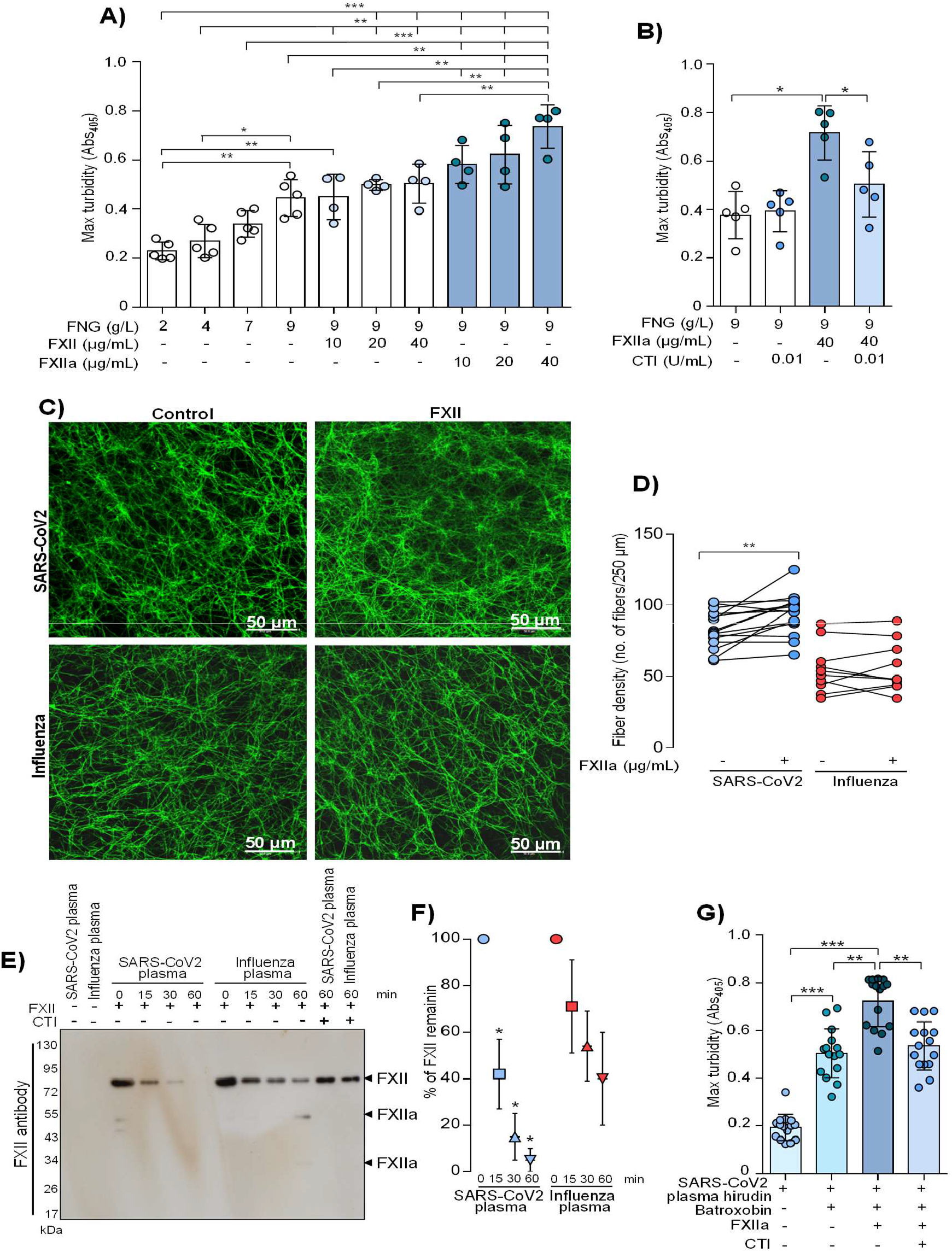
Fibrinogen and FXIIa contribute to dense fibrin network in severe COVID-19. A, B) Max turbidity values of fibrin clots generated in the purified system from increasing concentrations of FNG and/or FXII/FXIIa in the absence or presence of CTI. Clot formation was induced by thrombin. n=4-5. C) Laser scanning confocal microscopy images of fibrin fibers in clots formed from SARS-CoV2 or Influenza plasma supplemented with factor XII (FXII). Representative pictures are demonstrated. D) Fibrin fiber density in ARDS-Influenza (n=10) and COVID-19 (n=10) clots generated in C). Per patient 3 separate clots were prepared, 5 pictures were taken in different areas of the clots and fibril density was determined in all pictures. Paired data is shown interconnected. E) Rate of FXII activation in ARDS-Influenza and SARS-CoV2 plasma. Biotin-labeled FXII was added to plasma and its decay was monitored by western blotting using horseradish peroxidase-labeled streptavidine (upper panel). Representative blot is shown. F) Quantification of FXII decay in ARDS-Influenza and SARS-CoV2 plasma. FXII signal at time point 0 was considered as 100%. n=20/group. G) Maximum (Max) turbidity values of fibrin clots generated by the addition of batroxobin to hirudin-preincubated plasma in the presence of active FXII (FXIIa) and/or corn trypsin inhibitor (CTI). n=15 biological replicates. Data in A), B), F) and (G) indicate mean ± SD. *p<0.05, **p<0.01, ***p<0.001.

As sustained activation of FXII was described in COVID-19^35^, we next investigated the potential contribution of FXIIa to the regulation of fibrin clot structure in severe COVID-19. Supplementation of COVID-19 plasma with FXII to the levels observed in healthy subjects increased fibrin network density but not fibrin fiber diameter. No apparent effect of FXII addition on fibrin clot architecture was seen in ARDS-influenza samples (Figure 3C, D). Furthermore, rapid decay of exogenous, biotinylated FXII in COVID-19, but not in ARDS-influenza, plasma, implying an accelerated rate of FXII activation in the former group of the patients was observed (Figure 3E, F). The addition of CTI to plasma samples prevented the conversion of FXII into FXIIa (Figure 3F). To demonstrate a direct effect of FXIIa on fibrin structure, we clotted hirudin-preincubated COVID-19 plasma with batroxobin in the presence of FXIIa and/or CTI and measured the maximum turbidity. As shown in figure 3G, FXIIa increased fibrin density and this effect was diminished by CTI thus ensuring direct, thrombin independent, role of proteolytically active FXII in the modulation of fibrin architecture. The elevated levels of factor XIIIa were not detected in plasma of critically ill COVID-19 patients (data not shown).

Together, these results suggest that high levels of fibrinogen along with markedly increased rates of FXII activation in COVID-19 plasma create a specific pro-coagulatory microenvironment, which promotes the formation of particularly dense fibrin networks.

### Elevated fibrin network density increases clot resistance to fibrinolysis

Fibrin network density was previously found to determine clot resistance to fibrinolysis^37^. Accordingly, we next evaluated the lysis resistance of fibrin clots in patient plasma, using an *in vitro* turbidimetric clot-lysis assay. Here, kaolin together with tissue-plasminogen activator (t-PA) were added to plasma to initiate the intrinsic pathway of coagulation, followed by fibrin-dependent plasmin generation *via* t-PA-mediated activation of plasminogen in the same sample. While in normal plasma, the characteristic bell-shaped clot-lysis curve, representing the complete fibrin clot dissolution, was observed, only partial clot-lysis was detected in ARDS-influenza samples, and clot-lysis was completely absent in COVID-19 samples over the entire time period of the experiment (Figure 4A). This observation is supported by the highest turbidity values at 60 min in severe COVID-19 samples (Figure 4B). Overall, clot lysis was observed in 84% of ARDS-influenza patients and only 30% of COVID-19 patients suggesting fibrinolysis shutdown in the vast majority of SARS-CoV2-infected patients in our cohort.

**Figure 4.**
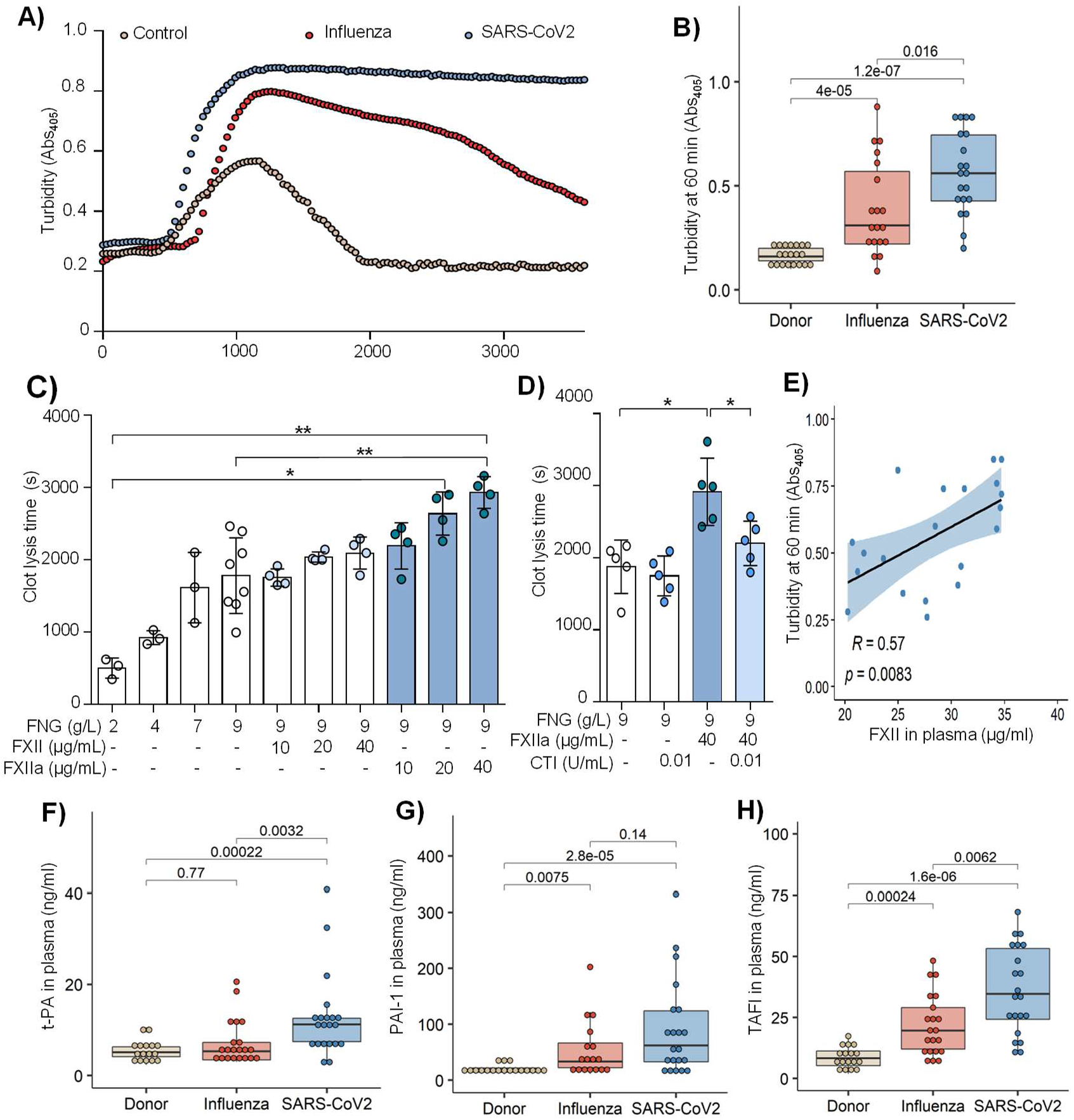
Fibrinolysis shutdown in severe COVID-19. A) Turbidimetric analysis of clot lysis in severe COVID-19 (SARS-CoV2), ARDS-influenza and control plasma. Representative clot lysis curves are shown. SARS-CoV2, n=20; ARDS-Influenza; n=19, control, n=20. B) Turbidity values (A_405_) of the fibrin clots at 60 min. SARS-CoV2, n=20; ARDS-influenza, n=19; control, n=20. C, D) Clot lysis time. Clots were generated in purified system with increasing concentrations of fibrinogen (FNG) and/or factor XII (FXII)/active FXII (FXIIa). Clot formation was induced by thrombin and clot lysis by plasmin generated from plasminogen by t-PA. In some experiments FXII was preincubated with corn trypsin inhibitor (CTI). Clot formation and lysis were monitored via turbidimetry. n=3-5. Mean+/-SD is shown. *p<0.05, **p<0.01, ***p<0.001. E) Correlation between turbidity values at 60 min and FXII plasma levels in severe COVID-19. n=20. Correlation is performed using Spearman’s rank correlation coefficient. F-H) t-PA (F), plasminogen activator inhibitor-1 (PAI-1; G), and thrombin-activatable fibrinolysis inhibitor (TAFI, H) levels in plasma of severe COVID-19 (n=21), ARDS-influenza (n=21) and control (n=17) as assessed by ELISA. Data in B) and F)-H) are shown as single data points with boxplot overlay indicating median and interquartile range.

To probe for high fibrinogen levels and increased rate of FXII activation as potential cause of fibrinolysis shutdown in plasma from critically ill COVID-19 patients, clot-lysis assays were performed in a purified system. As expected, increasing amounts of fibrinogen and FXIIa prolonged clot lysis time, with an additive effect being observed at the highest concentrations of both proteins (Figure 4C). The addition of CTI to the assay shortened clot lysis time supporting the requirement of FXII proteolytic activity for this effect (Figure 4D). In addition, a strong positive correlation between the turbidity values at 60 min and FXII levels in COVID-19 plasma was seen (Figure 4E).

To test whether other components of the fibrinolytic system, such as t-PA, plasminogen activator inhibitor-1 (PAI-1) and thrombin-activatable fibrinolysis inhibitor (TAFI, also designated plasma carboxypeptidase B2) may be dysregulated in critically ill COVID-19 patients, we measured their levels by means of ELISA. The concentration of t-PA was elevated in severe COVID-19 as compared to control and ARDS-influenza patients (Figure 4F). An increase of PAI-1 was also noted in plasma of ARDS-influenza and severe COVID-19 patients as opposed to control, yet, a significant difference between both patient groups was not detected (Figure 4G). Interestingly, TAFI was not only markedly elevated in both patient groups as compared to control, but exhibited also significantly higher plasma values in severely ill patients with COVID-19 as compared to ARDS-influenza (Figure 4H).

### Dense fibrin clots are observed in the lungs of severe COVID-19 patients

To demonstrate the *in vivo* relevance of our findings, we stained autopsy lung tissue sections from SARS-CoV2- and influenza-infected ARDS patients as well as subjects who died due to no respiratory causes for fibrin. Notably, time from death to autopsy was matched for all groups examined. As demonstrated in figure 5A, intra- and extra-vascular fibrin aggregates were observed in both severe COVID-19 and ARDS-influenza patients. However, in contrast to ARDS-influenza subjects, in the lungs of COVID-19 patients the deposits of fibrin appeared to be more widespread and evenly present not only in alveolar spaces but also around alveolar septae over the whole lung examined. In ARDS-influenza patients, fibrin deposit were predominantly observed in alveolar spaces and present in selected regions of the lung (Figure 5A). Overall, in COVID-19 lungs fibrin clots were more compact and homogeneous whereas in ARDS-influenza lungs they were widespread and characterized by regions of high and low fibrin fiber density (Figure 5B).

**Figure 5.**
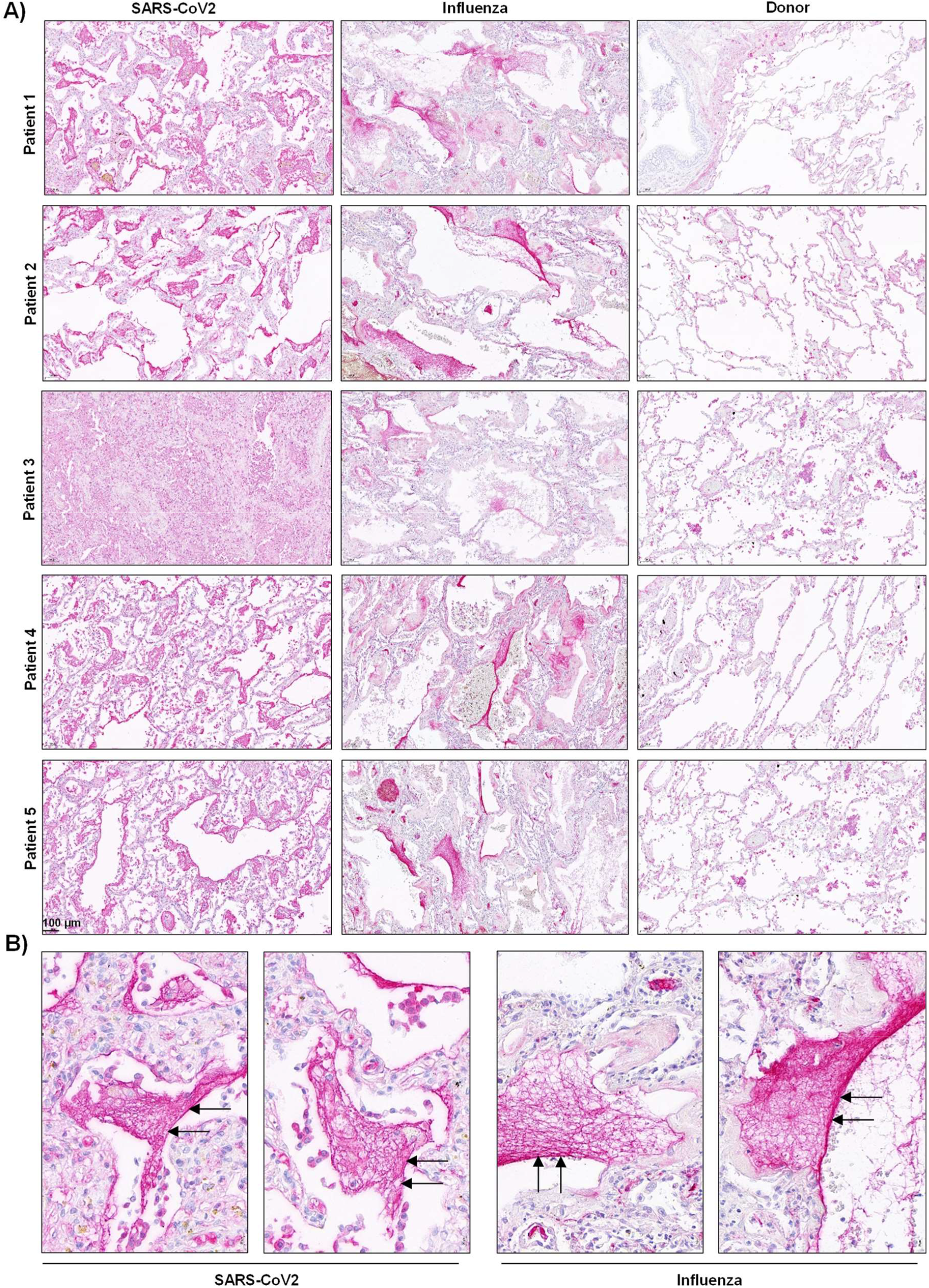
Fibrin deposits in the lungs of severe COVID-19 and ARDS-influenza patients. A, B) Fibrin (red) accumulation in postmortem lung tissue sections of severe COVID-19 (n=5), ARDS-influenza (n=5) and donors (n=5). Time from death to autopsy was matched for all groups examined. Arrows indicate fibrin deposits in the lung. Magnification bar 100 µm.

## Discussion

Many patients with severe COVID-19 exhibit coagulation abnormalities that mimic other systemic coagulopathies associated with severe infections, such as disseminated intravascular coagulation (DIC) or thrombotic microangiopathy^36^. A high incidence of venous thromboembolism, pulmonary embolism, deep vein thrombosis, and multiple organ failure with a poor prognosis and outcome appears to be causally related to dysregulation of blood coagulation in critically ill COVID-19 patients. Besides an elevated inflammatory status (e.g. increased cytokine levels) that might induce monocyte-related coagulation and suppression of anticoagulant pathways, typical laboratory findings in COVID-19 patients with coagulopathy are increased D-dimer levels and elevated fibrinogen concentrations^36^. Moreover, inflammation-induced endothelial cell injury in different vascular beds may contribute to a hypercoagulable state and the risk of thromboembolic complications^37,38^.

In order to provide mechanistic insights into the reported hypercoagulable state of severe COVID-19 patients, we compared changes in the contact phase system activation and fibrinolysis between COVID-19 patients, individuals suffering from ARDS-influenza, and donors. While some critical parameters such as fibrinogen, PAI-1, and TAFI were significantly increased, FXII levels were reduced in severe COVID-19, and the process of fibrin formation and the resulting fibrin clot structure and lysis were substantially different between patient cohorts. Histological data provided evidence for widespread, compact fibrin deposition in the lungs of patients with COVID-19 as opposed to those with ARDS-influenza.

In particular, although the levels of FXII were significantly decreased in severe COVID-19 patients as compared to ARDS-influenza and donors, FXII-activation products were markedly altered in patients with SARS-CoV2 infection. This scenario very likely reflects FXII consumption due to its increased binding to and auto-activation on negatively charged surfaces. Decreased FXII levels in COVID-19 plasma are also in accordance with moderately elevated APPT reported in other studies^39,40^. The exacerbated consumption of FXII in severe COVID-19 is further supported by our *in vitro* studies, in which the supplementation of COVID-19 plasma with exogenous FXII resulted in its rapid activation, presumably due to the presence of FXII auto-activation cofactors. Indeed, common pathological events observed in COVID-19 such as increased tissue cell stress together with virus-mediated necrosis, endothelial dysfunction, and excessive neutrophil activation, lead to the release/exposure of large amounts of negatively charged molecules including NETs. NETs not only bind FXII but also serve as a potent endogenous inflammation-dependent inducer of FXII auto-activation, eventually propagating thrombosis^21,41^. Enhanced vascular NETosis along with impaired NET clearance were described in COVID-19 patients^35,42^. In line with these findings, several studies found an increase in NET components in COVID-19 plasma including cell-free DNA, myeloperoxidase-DNA complexes, neutrophil elastase-DNA complexes, and citrullinated histone H3^43,44^. In addition, active FXII was described to colocalized with NETs in the lungs of COVID-19 patients and NET positive pulmonary vessels were reported to be frequently clogged^38,45^. Together with these findings, our results speak for NET-induced, accelerated, and constant activation of FXII in COVID-19 and thus for its role in immunothrombothic processes in this pathology. In fact, FXII auto-activation cofactors were found to be relevant for the initiation and progression of sepsis and DIC^46^.

Interestingly enough, low plasma levels of FXII in severe COVID-19 patients did not result in markedly prolonged kaolin clotting time (KCT) suggesting that other hemostatic abnormalities/factors compensate for low amounts of FXII in critically ill COVID-19 subjects. As previous studies reported that high plasma levels of FVIII:C may associate with a short KCT and an increased risk of thromboembolism^47^, it is plausible to assume that the excessive amounts of FVIII:C in COVID-19, as opposed to ARDS-influenza, plasma induced shortening of KCT in our cohort of patients. These results, together with previously described high levels of fibrinogen, mild thrombocytopenia, and slightly altered plasma concentrations of coagulation factors and physiological anticoagulants^48^ argue for a specific form of intravascular coagulation in severe COVID-19 that is distinguishable from classical DIC. The prominent increase in vascular complications^41^ points to strong involvement of endothelial cells in hemostatic abnormalities seen in COVID-19. Injured endothelial cells may provide a scaffold for thrombus generation and elevated levels of von Willebrand factor multimers (recently described in COVID-19 plasma^49^) may facilitate platelet-vessel wall interactions ultimately leading to the formation of platelet-rich thrombotic deposits in microvasculature. Such platelet-rich thrombotic aggregates have been observed in alveolar capillaries of critically ill COVID-19 patients^16,17^. Altogether haemostatic alterations seen in COVID-19 subjects reflect widespread occlusive thrombotic microangiopathy with destruction of alveoli that supports persistence of microthrombi.

Elevated levels of fibrinogen were reported to contribute to the faster fibrin formation and increased fibrin network density, strength, and stability^34^. In line with this assumption, clots generated from COVID-19 plasma exhibited much higher packing density as compared to those formed from ARDS-influenza plasma. Further experiments with COVID-19 plasma and in a purified system revealed that next to fibrinogen also FXIIa may regulate clot compactness. Indeed, higher levels of fibrinogen and increased rate of FXII activation were associated with denser fibrin clots with smaller pores. The compact architecture of clots generated from COVID-19 plasma correlated with their resistance to lysis consolidating the notion of hyperfibrinogenemia and FXII consumption coagulopathy as driving causes of an increased risk of thrombosis in critically ill COVID-19 patients. Our findings are consistent with the studies demonstrating the role of fibrinogen and FXIIa in organization of clot architecture^33,34^ and the reports linking abnormal fibrin network structure/function with thrombotic events seen in patients with diabetes^50^, ischemic stroke^51^, pulmonary hypertension^52^, myocardial infraction^53^, or venous thromboembolism^54^. Although, increased fibrinogen levels independently promote thrombus formation and stability, the role of FXII in these processes seems to be more complex and dependent on environment conditions. Those include, the presence of NETs (or any other molecule being able to activate FXII) which orchestrate not only FXII but also platelets activation, activated platelets may perpetuate FXIIa generation by the release of polyphosphates and the availability of haemostatic factors. Coagulation proteases ensure FXIIa-dependent thrombin formation and a direct binding of FXII/FXIIa to fibrinogen may define aggregation of fibrin fibers^33^. Whether the interaction of FXII/FXIIa with fibrinogen can interfere with the binding of t-PA to fibrin and thereby inhibits fibrinolysis warrants further investigation.

Clots generated from COVID-19 plasma exhibited higher packing density, small pores and were built of thin fibers. Interestingly enough, previous studies suggested that thrombi made of thin and numerous fibers organized in tight network are resistant to fibrinolysis^55^. Persistent vessel occlusion seen in critically ill COVID-19 patients is reinforced by markedly increased plasma levels of TAFI and moderately elevated amounts of PAI-1^56^. Thus, persistent occlusion of microvessels in the lungs of COVID-19 patients appears to be a result of unfortunate circumstances, starting from sustained activation/presence of thrombosis-promoting factors, going through the formation of lysis resistant thrombi, and finishing on the accumulation of fibrinolytic inhibitors^57^.

Based on current and previous findings, the scenario of defense mechanisms, including the immune and coagulation system, running out of control emerges as an underlying mechanism for severe SARS-CoV2 infection. Multiple hits from abnormalities in plasma composition, vascular cell function, and blood immune cell landscape through virus-mediated cell damage and release of intracellular debris create a milieu favoring activation of FXII. In combination with high levels of fibrinogen, FXIIa contributes to pathologic thrombus formation not only *via* thrombin generation but also through the formation of compact and lysis resistant clots. Our study thus establishes a model for future investigations on the role of altered fibrin clot structure in thrombosis and thrombolysis in severe COVID-19.

## Materials and Methods

### Study population

Plasma samples from COVID-19 patients were obtained from the Hannover Medical School, Hannover, Germany (validation cohort) and from the Charité-University Medicine, Berlin, Germany (discovery cohort). Plasma samples from acute respiratory distress syndrome (ARDS) due to influenza were provided from the Hannover Medical School, Hannover, Germany. All samples were taken within 6 days after onset of ARDS. All investigations were approved by the local ethics committees (Hanover samples: SEPSIS/ARDS Registry, ethic votum no.: 8146_BO_K_2018; Berlin samples: ethic votum no.: EA2/066/20) and written informed consent was obtained from all participants or their next-of-kin. COVID-19 patients were classified as moderate (hospitalized, no invasive ventilation; WHO severity score: 3-4) or severe (high flow O_2_ or intubated and mechanically ventilated; WHO severity score: 5-7) as previously described^58^. Control (healthy subjects) samples were provided by the Charité-University Medicine, Berlin, Germany (ethic votum no.: EA2/075/15) and from the Justus-Liebig University of Giessen, Giessen, Germany (ethic votum no.: 05/00). Baseline demographics and clinical characteristics of the patients are shown in Table 1.

**Table 1.** Baseline demographics and clinical characteristics of COVID-19 and ARDS-influenza patients (plasma samples).

|  | Clotting |  |  |  |
| --- | --- | --- | --- | --- |
|  | ARDS<sup>1</sup>-influenza | COVID<sup>2</sup>-19<br>(WHO 5-7) | COVID-19<br>(WHO 5-7) | COVID-19<br>(WHO 3-4) |
| No. of patients, n | 20 | 21 | 15 | 15 |
| Age, year | 56; [20-86] | 59; [19-82] | 61; [22-84] | 61; [26-80] |
| Sex, male (%) | 87 | 90 | 69 | 67 |
| BMI (kg/m<sup>2</sup>) | 25; [20-36] | 29; [15-62] | 29; [25-36] | 24; [20-36] |
| <sup>3</sup>CRB65 score, n |  |  |  |  |
| 0 | 0 | 0 | NA | NA |
| 1 | 0 | 2 |  |  |
| 2 | 3 | 6 |  |  |
| 3 | 12 | 9 |  |  |
| 4 | 5 | 4 |  |  |
| 28-day mortality (%) | 30 | 14.3 | 8 | 0 |
| <sup>4</sup>LOS ICU, days | 19; [6-73] | 27; [3-63] | 30; [5-220] | NA |
| Ventilation, days | 15; [3-66] | 16; [4-50] | 26; [5-220] | NA |
| <sup>5</sup>ECMO (%) | 30 | 24 | 46 | NA |
| <sup>6</sup>SOFA | 10; [5-16] | 13; [9-17] | 10; [2-12] | NA |
| <sup>4</sup>CRP (mg/L) | 264 [31-406] | 151; [68-292] | 85; [27-411] | 29; [1-148] |
| Leukocytes, ×10<sup>9</sup>/L | 16; [22-90] | 9; [4-36] | 10; [5-27] | 7; [4-22] |
| Platelet count, ×10<sup>9</sup>/L | 199; [70-653] | 247; [99-581] | 286; [129-635] | 334; [173-602] |
| Lactate, mM | 1.3; [0.7-4.8] | 1.8; [0.7-5.6] | 1.7; [0.4-6.6] | 1.3; [1.0-2.6] |
| Procalcitonin, µg/L | 1.7; [0.2-79.5] | 0.6; [0.1-66.1] | 0.6; [0.1-25] | 0.1; [0-1] |
| D-dimer, mg/L | NA | 4; [1-35] | NA | NA |
| Fibrinogen, g/L | 4.0 [3.2-9.0] | 8.0; [4.0-9.0] | NA | NA |
| Anticoagulant |  |  |  |  |
| Heparin, n | 15 | 14 | 8 | 7 |
| <sup>8</sup>PTT, s | 38; [25-59] | 36; [26-55] | 41; [30-67] | 34; [30-60] |

<sup>1</sup>ARDS, acute respiratory distress syndrome; <sup>2</sup>COVID-19, coronavirus disease 2019; <sup>3</sup>CRB 65, confusion, respiratory rate, blood pressure, age 65 score; <sup>4</sup>LOS ICU, length of intensive care unit stay; <sup>5</sup>ECMO, extracorporeal membrane oxygenation; <sup>6</sup>SOFA, sequential organ failure assessment; <sup>7</sup>CRP, C-reactive protein; <sup>8</sup>PTT, partial thromboplastin time.

Lung specimens were obtained from 8 ARDS patients (5 COVID-19, 3 influenza) and 5 donors by autopsy. Time from death to autopsy was matched for all groups. All investigations were approved by the local ethics committees (Medical Faculty of Justus-Liebig University of Giessen, ethic votum no.: 29/01 and Medical University of Graz, ethic votum no.: 32-362 ex 19/20) and written informed consent was obtained from all participants or their next-of-kin if required. Baseline demographics and clinical characteristics of lung tissue donors are shown in Table 2.

**Table 2.** Baseline demographics and clinical characteristics of COVID-19 and ARDS-influenza patients (lung tissue).

| COVID-19 |  |  |  |  |  |
| --- | --- | --- | --- | --- | --- |
| Patient | Age, year | Sex | Background | Ventilation, days | Anticoagulant |
| 1 | 82 | Male | Diffuse alveolar damage | 0 | Heparin |
| 2 | 77 | Male | Diffuse alveolar damage | 2 | Heparin |
| 3 | 72 | Male | Diffuse alveolar damage | 6 | Heparin |
| 4 | 65 | Male | Diffuse alveolar damage | 33 | Heparin |
| 5 | 79 | Female | Diffuse alveolar damage | 2 | Heparin |
| ARDS-influenza |  |  |  |  |  |
| 1 | 67 | Male | Community acquired pneumonia | 6 | Heparin |
| 2 | 72 | Male | Trauma | 3 | Heparin |
| 3 | 77 | Male | Community acquired pneumonia | 5 | Heparin |
| 4 | 81 | Female | Community acquired pneumonia | 10 | Heparin |
| 5 | 80 | Female | Trauma | 2 | Heparin |
| Donor |  |  |  |  |  |
| 1 | 82 | Female | Recurrent myocardial infarction | 0 | no |
| 2 | 75 | Female | Heart and lung failure | 0 | Heparin |
| 3 | 64 | Male | Myocardial infarction | 0 | Heparin |
| 4 | 77 | Female | Dilated cardiomyopathy (right) | 0 | Heparin |
| 5 | 75 | Male | Dilated cardiomyopathy (right) | 0 | Heparin |

<sup>1</sup>ARDS, acute respiratory distress syndrome; <sup>2</sup>COVID-19, coronavirus disease 2019.

### Plasma clot formation and lysis

Twenty μL of plasma were preincubated for 10 min with 20 μL of 0.1 M imidazole buffer, pH 7.4, and 20 μL of 0.3 mg/mL kaolin in a clear, flat-bottomed 96-well plate. Clotting was initiated by the addition of 20 μL of 20 mM CaCl_2_ in the absence or presence of tissue plasminogen activator (t-PA) (25 ng/mL final; Sekisui Diagnostics, Burlington, MA). Turbidity was monitored at 405 nm (A_405_) every 30 s for 60 min at 37°C using a SpectraMax 190 (Molecular Devices, Biberach, Germany). In some experiments, COVID-19 plasma was preincubated with hirudin (5 IE/mL final; Diapharma, West Chester, OH) and the clotting was induced by batroxobin (5U/mL final; Enzyme Research Laboratories, South Bend, IN)

### Fibrin formation and lysis in a purified system

Thrombin (5 nM final, Sekisui Diagnostics) was mixed with fibrinogen (2-9 g/L final), pre-incubated with either FXII or FXIIa (10-40 μg/mL final, both from Sekisui Diagnostics) in a total volume of 25 μL of 0.1 M imidazole buffer in a clear, flat-bottomed 96-well plate. Fibrin formation was initiated by the addition of 20 μL of 20 mM CaCl_2_. To measure fibrinolysis, t-PA (0.1 µg/mL final) and plasminogen (20 µg/mL final, Enzyme Research Laboratories) were added to the clotting solution. Turbidity was monitored as described above. In some experiments, FXIIa (40 μg/mL final) was incubated with corn trypsin inhibitor (CTI; 0.01 U/mL final, Sekisui Diagnostics) before mixing with fibrinogen.

### Western blotting

Plasma (pre-diluted 1:40 into 0.9% NaCl) was separated on a SDS polyacrylamide gel, followed by electro-transfer to a PVDF membrane. After blocking with 5% non-fat dry milk in TBS buffer (25 mM Tris pH 7.5, 150 mM NaCl) supplemented with 0.1% Tween 20 (TBS-T), the membrane was incubated overnight at 4°C with a goat anti-FXII (cat. no.: 206-0056; Zytomed Systems, Berlin, Germany) or rabbit-anti high molecular weight kininogen (HK; cat. no.: ab35105; Abcam, Cambridge, UK) antibody. Next, membranes were incubated with appropriate peroxidase-labelled secondary antibodies (all from Dako, Gostrup, Denmark). Final detection of proteins was performed using a PierceTM ECL Western Blotting Substrate (Thermo-Fisher Scientific). As loading control, albumin was detected with a rabbit anti-albumin antibody (cat. no.: A001; Dako). Western blots were developed using a ChemiDocTM Touch (BioRad Laboratories, Inc., Hercules, CA), and densitometric analysis was conducted by the ImageLabTM, Version 6.0.1 (Bio-Rad Laboratories).

### Immunoassays

Factor XII levels in plasma were quantified by the Human FXII ELISA Kit from Abnova (Taipei, Taiwan). Plasma levels of plasminogen activator inhibitor-1 (PAI-1) and t-PA, were measured using human ELISA Kits from Thermo-Fisher Scientific. Thrombin-activatable fibrinolysis inhibitor (TAFI) levels in plasma were quantified by Human CPB2/TAFI ELISA Kit from LSBio (Seattle, WA). All measurements were performed according to manufacturer’s instructions.

### FXII decay in plasma

Endogenous FXII was depleted from plasma using a goat anti-FXII antibody (cat. no.: 206-0056; Zytomed Systems) covalently attached to magnetic beads (Thermo-Fisher Scientific). Afterwards, a hundred µl of plasma was supplemented with 1 nM biotinylated FXII and the sample was incubated for 1h at 37°C. Aliquots were withdrawn after the indicated time points and analyzed by western blotting. In some experiments, plasma was preincubated with 12 mg/mL CTI 30 min prior to the addition of biotinylated FXII.

### Immunostaining of clots generated in a purified system

Clots were generated from Influenza and Covid-19 plasma supplemented with 10 µg/ml exogenous FXII as described above. Next, they were fixed with 4% paraformaldehyde in PBS. Non-specific binding sites were blocked with 3% BSA in PBS for 1 h. Next, clots were incubated with a rabbit anti-fibrinogen/fibrin (cat. no.: A 0082; Dako) antibody overnight at 4°C. Following extensive washing with PBS, clots were incubated with secondary antibodies labeled with Alexa Fluor™ 488 (Thermo-Fisher Scientific) for 1h at room temperature. Finally, clots were embedded in Vectashield Mounting Medium (Vector Laboratories Inc) and images were taken as described above. ImageJ was used to determine fiber density, by counting the number of fibers crossing lines of 250 µm placed in the image using the plug-in-grid.

### Scanning electron microscopy

Samples were fixed with 1.5% paraformaldehyde and 1.5% glutaraldehyde solution in 0.15 M Hepes for 24 h at room temperature. Next, samples were washed with 0.15 M Hepes, post-fixed in 1% osmium tetroxide for 2 h, washed in distilled water, dehydrated with graded ethanol washes and critical point dried by CO_2_ treatment using a CPD 030 critical point dryer (Evatec AG, Trübbach, Switzerland). Finally, samples were mounted with conductive adhesive tape and sputtered with gold. Images were taken with a Philips XL30 scanning electron microscope (Philips, Eindhoven, Netherlands).

### Activity assays

The PKa-like activity assay and the activity of factor VIII were performed as described in ^59^ and ^60^, respectively.

### Statistics

Statistical analysis was performed in R (version 4) using the ggpubr package. Data are expressed as single data points with boxplot overlay indicating median and interquartile range, unless indicated otherwise. Multiple groups were compared by non-parametric Kruskal-Wallis test. Correlations were performed using Spearman’s rank correlation coefficient.

## Competing interests

None declared.

## Acknowledgement

We thank E. Bieniek for her excellent technical assistance. We also thanks A. Seipp (Institute of Anatomy and Cell Biology, Justus Liebig University, Giessen, Germany) for performing the scanning electron microscopy pictures. This study was funded by the German Research Foundation (DFG: SFB/TR84 Project A2 to M.W. and W.M.K.), the Else Kröner-Fresenius-Foundation (2014_A179 to M.W. and P.M.), the Oskar Helene Heim Foundation (to P.M.), the German Center for Lung Research (82 DZL 005A1 to M.W.) and the University Medical Center Giessen and Marburg (UKGM to M.W. and P.M.).

## Contribution

M.W. designed the study, performed experiments, analyzed data, and wrote the manuscript; A.B., L.M., and O.P. performed experiments and analyzed data; B.S., S.D., T.W., J.J.S., M.C.B., S.H., F.K., L.E.S., and M.Wi. recruited patients, analyzed patient clinical data, and reviewed the manuscript; A-S.S. and F.S. analyzed patient clinical data and wrote the manuscript; M.Z. and G.G. collected autopsy tissue samples and reviewed the manuscript; N.W., R.T.S., G.B., L.S., and P.M. analyzed data and contributed to the writing of the manuscript; W.M.K., G.K., and K.T.P designed the study and wrote the manuscript.

## Notes

### Competing Interest Statement

The authors have declared no competing interest.

